# Low-dose bivalent mRNA vaccine is highly effective against different SARS-CoV-2 variants in a transgenic mouse model

**DOI:** 10.1101/2022.04.20.485440

**Authors:** Björn Corleis, Donata Hoffmann, Susanne Rauch, Charlie Fricke, Nicole Roth, Janina Gergen, Kristina Kovacikova, Kore Schlottau, Nico Joel Halwe, Lorenz Ulrich, Jacob Schön, Kerstin Wernike, Marek Widera, Sandra Ciesek, Stefan O. Mueller, Thomas C. Mettenleiter, Benjamin Petsch, Martin Beer, Anca Dorhoi

**Author notes:** **Contributor Information:** Contributed equally (Björn Corleis, Donata Hoffmann, Martin Beer, Anca Dorhoi).

## Abstract

Combining optimized spike (S) protein-encoding mRNA vaccines to target multiple SARS-CoV-2 variants could improve COVID-19 control. We compared monovalent and bivalent mRNA vaccines encoding B.1.351 (Beta) and/or B.1.617.2 (Delta) SARS-CoV-2 S-protein, primarily in a transgenic mouse model and a Wistar rat model. The low-dose bivalent mRNA vaccine contained half the mRNA of each respective monovalent vaccine, but induced comparable neutralizing antibody titres, enrichment of lung-resident memory CD8^+^ T cells, specific CD4^+^ and CD8^+^ responses, and fully protected transgenic mice from SARS-CoV-2 lethality. The bivalent mRNA vaccine significantly reduced viral replication in both Beta- and Delta-challenged mice. Sera from bivalent mRNA vaccine immunized Wistar rats also contained neutralizing antibodies against the B.1.1.529 (Omicron BA.1) variant. These data suggest that low-dose and fit-for-purpose multivalent mRNA vaccines encoding distinct S-proteins is a feasible approach for increasing the potency of vaccines against emerging and co-circulating SARS-CoV-2 variants.

## Main text

Effective vaccines are critical for the control of the COVID-19 pandemic, especially as nations begin to scale back non-pharmaceutical interventions such as social distancing, travel restrictions, and isolation (https://www.ecdc.europa.eu/en/covid-19/prevention-and-control/vaccines).

Since the beginning of the pandemic, new SARS-CoV-2 variants, including some classed as variants of concern (VOCs) have appeared, each characterized by different virulence, transmissibility, and immune escape resulting in differences in the effectiveness of public health measures, diagnostics, vaccines, or therapeutics. While B.1.1.7 (Alpha) and B.1.617.2 (Delta) spread rapidly, particularly in the naïve population, B.1.351 (Beta) and especially B.1.1.529 (Omicron) are notable for immune escape (https://www.who.int/en/activities/tracking-SARS-CoV-2-variants).

Several VOCs which have mutated extensively, such as Beta^1^ and Omicron,^2^ have been able to evade humoral responses elicited by vaccines based on ancestral S-protein sequences.^3^ As a result, Omicron has quickly become globally prevalent, despite high immunization rates. Unfortunately, while Omicron appears to cause less severe disease than other variants,^4^ it does not induce relevant cross-protective neutralizing antibody (nAb) titres in SARS-CoV-2 naïve populations, meaning they may be less protected against future infection compared with those previously exposed to other variants or vaccinated.^5^

The evolution of further VOCs is unpredictable; however, it is likely that new escape variants will emerge. Therefore, developing effective vaccines and vaccine strategies will remain essential.^6^ SARS-CoV-2 vaccine development should use proven concepts of licensed vaccines to aid optimization (https://www.ema.europa.eu/en/human-regulatory/overview/public-health-threats/coronavirus-disease-covid-19/treatments-vaccines/vaccines-covid-19/covid-19-vaccines-authorised), with broad protection against different VOCs. For example, prototypes of multivalent nanoparticle vaccines have been reported to induce broad reactivity against different sarbecoviruses.^7^

mRNA vaccines are promising candidates for future vaccine approaches based on their demonstrated ability to induce robust protection against SARS-CoV-2.^8^ The development of low-dose multivalent human vaccine preparations against different VOCs is an innovative approach. We designed unmodified mRNA vaccines encoding the S-protein sequences from the Beta and Delta variants, which are distant variants with non-overlapping mutations in the RBD domain, as well as the ancestral strain; CV2CoV.351, CV2CoV.617 and CV2CoV, respectively.^2,9^ The vaccines containing a total of 0.5 μg mRNA per dose were administered intramuscularly; the bivalent vaccine (CV2CoV.351 and CV2CoV.617 together) contained 0.25 μg mRNA of each variant, i.e. half the dose of the monovalent vaccines.

K18-hACE2 transgenic mice received 20 μL of a low dose^10^ monovalent CV2CoV, CV2CoV.351, CV2CoV.617 or bivalent vaccine (CV2CoV.351 and CV2CoV.617) containing a total of 0.5 μg mRNA or NaCl (sham) on Day 0 and Day 28 (Fig. S1). These transgenic mice have a human ACE2 receptor which is the major cell entry receptor for SARS-CoV-2.^11,12^ Following challenge with Beta and Delta variants (10^4,4^ TCID_50_) on Day 56, all vaccinated mice were protected from SARS-CoV-2-induced lethality and virus spread, while all Beta-challenged and 67% of Delta-challenged sham-vaccinated animals succumbed to infection (Fig. 1a,b and Fig. S2a,b).

**Fig. 1.**
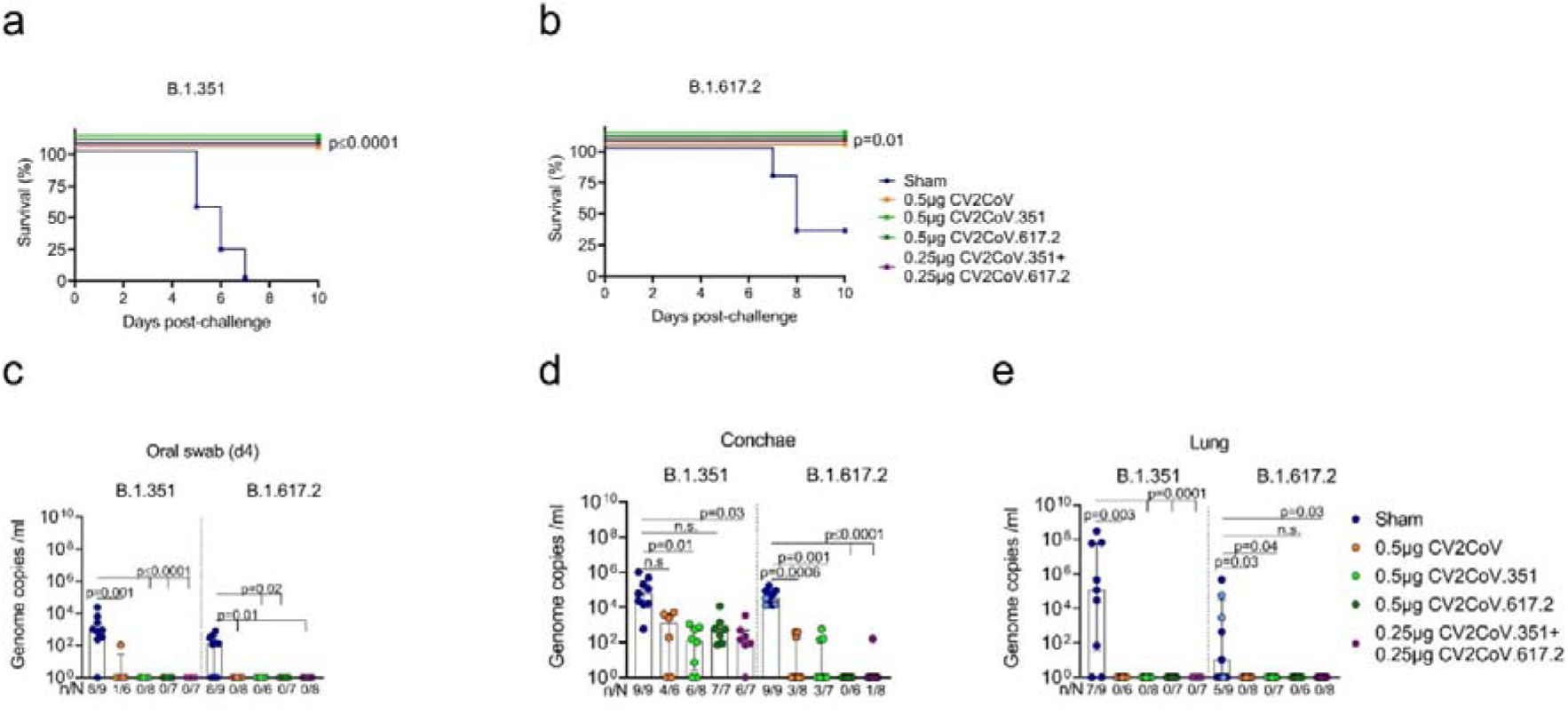
Monovalent and bivalent mRNA vaccines encoding ancestral, Beta and Delta derived S-protein sequences protect against SARS-CoV-2 variants in transgenic mouse model. K18-hACE2 mice vaccinated on day 0 and 28 with a total of 0.5 μg CV2CoV (ancestral, orange), 0.5 μg CV2CoV.351 (Beta, light green), 0.5 μg CV2CoV.617.2 (Delta, dark green), CV2CoV.351 + CV2CoV.617.2 (0.25 μg of each; purple) or NaCl (sham; blue) were challenged i.n. with 10^4,4^ TCID_50_ SARS-CoV-2 variant B.1.351 (Beta) or B.1.617.2 (Delta) at day 56. **(a,b)** Survival curves (Kaplan-Meier) for K18-hACE2 mice challenged with B.1.351 (Beta) **(a)** or B.1.617.2 (Delta) **(b)** with follow-up for 10 days post challenge. **(c-g)** RT-qPCR results from Day 4 oral swabs **(c)** or Day 10 conchae **(d)** and lung **(e).** Sham group samples were obtained at Day 10 (light blue) or at the humane endpoint (dark blue). Number of RT-qPCR positive and total number of animal sample are shown on the x-axis. Each dot represents one individual mouse. Scatter plots are labelled with median and interquartile range. *p-*values were determined by two-sided log-rank (Mantel-Cox) test **(a,b)** or one-way ANOVA and Dunn’s multiple comparison test **(c-e)**. Differences were considered significant at *p* □ 0.05 with exact *p* values shown.

Neither SARS-CoV-2 genomic RNA (Fig. 1c-e and Fig. S2c,d) or subgenomic RNA (sgRNA) (Table S2 and S3) were detectable in oral swabs collected on Day 4, or in lung, cerebellum and cerebrum samples taken on Day 10 in all but one of the Beta-challenged, bivalent-vaccinated animals, indicating that productive infection was prevented. The suppression of viral replication in the upper respiratory tract (URT) by the monovalent vaccines differed depending on the challenge virus. While the bivalent vaccine reduced viral load in the conchae equivalent to that observed with the matched monovalent vaccine after Beta challenge (Fig. 1d), replication of the Delta variant in the conchae was abolished (no detectable sgRNA) in all vaccinated groups (Fig. 1d and Table S3). Recent studies have revealed that repeated immunization extends neutralization to non-homologous variants, possibly through affinity maturation of memory B cell populations.^9,13^ Following the effective virus clearance in conchae, and the subsequent impact on Delta transmission, it will be important to evaluate the impact of bivalent, and higher valency, mRNA vaccines on nonneutralizing antibody functions, e.g., antibody-dependent natural killer cell activation or cellular phagocytosis, which are elicited by mRNA vaccines and maintained despite reductions in nAbs over time.^14^ These non-neutralizing antibodies may facilitate virus clearance like that observed in the URT of Delta-challenged mice.

Multivalent influenza^15^ and cytomegalovirus^16^ mRNA vaccines in pre-clinical models have shown that integration of multiple antigens can lead to robust nAb responses. Furthermore, nAbs have emerged as correlates of immune protection and vaccine efficacy^17^ to inform immunization schedules.^18^ In our study, anti-RBD total immunoglobulin levels were high in all vaccinated mice, with no notable differences between groups (Fig. S2e,f). The bivalent mRNA vaccine induced similarly high nAb titres as the Beta and Delta monovalent vaccines with their respective homologous challenges (Fig. 2a,b), despite the bivalent vaccine containing half the mRNA dose of each monovalent vaccine (0.25 μg vs. 0.5 μg). Irrespective of the challenge, the bivalent mRNA vaccine-induced nAb titres were statistically significantly higher than those induced by CV2CoV, whereas with the monovalent mRNA vaccines the nAB titres were statistically significantly higher with the respective homologous challenges only (Fig. 2a,b).

**Fig. 2.**
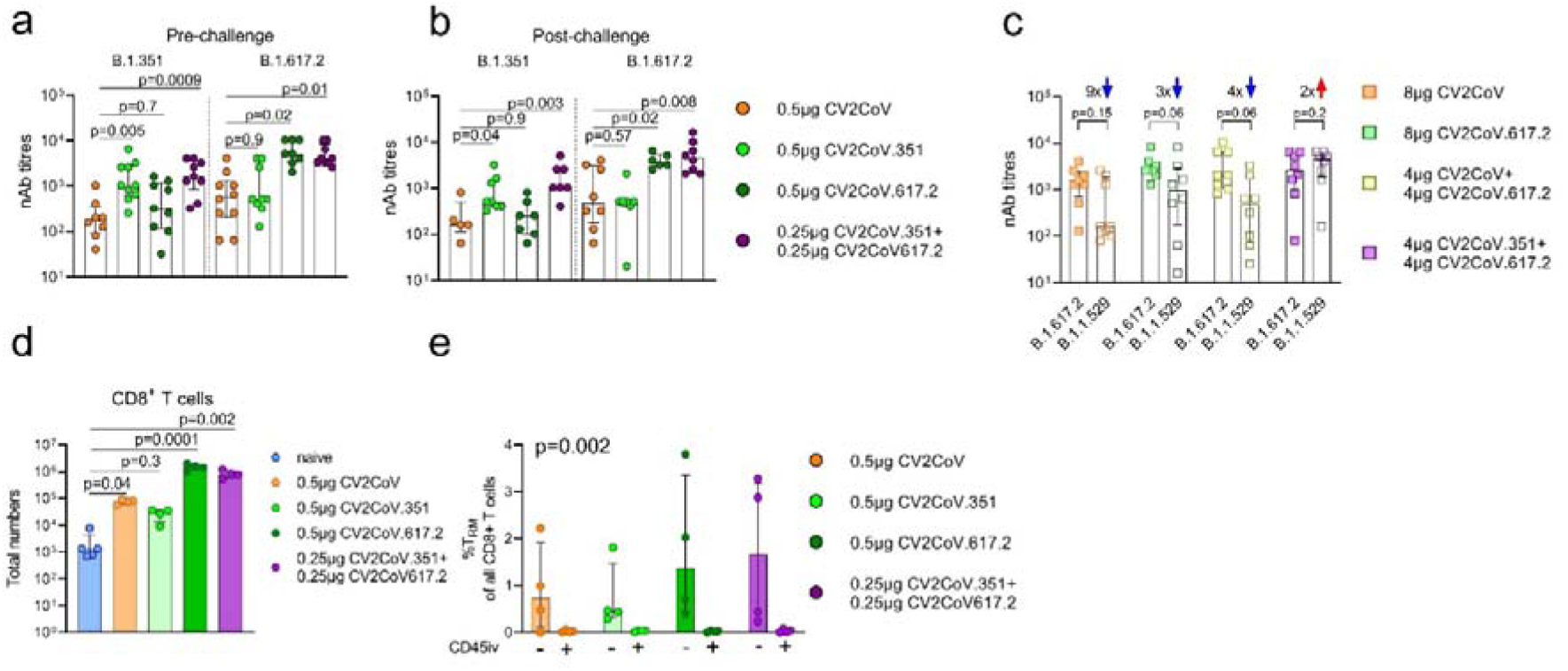
Bivalent SARS-CoV-2 mRNA vaccines induce abundant virus neutralizing titres and robust lung T cell responses. **(a,b)** Neutralizing antibody (nAb) titres (at Day 56 (Pre-challenge, **a**) and Day 66 (Postchallenge, **b**). Mice were vaccinated as described in Fig. 1. **(c)** nAb titres at Day 42 (prechallenge; Wistar rats), the numbers above the bars indicate the fold-differences. Rats were vaccinated on day 0 and 21 as described in Fig. S5 and nAb titres against B. 1.617.2 (Delta) or B.1.1.529 (Omicron, BA. 1) were tested. Each dot represents an individual animal. **(d-e)** Induction of low-dose mRNA vaccine induced T cell responses. Lung parenchyma T cells at Day 56 post vaccination were investigated by *in vivo* injection of 3 μg anti-mouse CD45 antibodies (CD45iv) for 3 minutes before harvesting of lung tissue. **(d)** Total number of CD45iv^−^ CD8^+^CD3^+^ T cells in all vaccine groups compared to naïve mice. **(e)** Frequency of CD8^+^ T_RM_ cells after vaccination and defined as CD45iv^−^CD3^+^γδTCR^−^CD8^+^CD44^high^CD62L^−^ CD103^+^CD69^+^ T cells. Scatter plots are labelled with median and interquartile range. *p-*values were determined by one-way ANOVA and Dunn’s multiple comparison test against CV2CoV (orange) **(a-c)** or against a group of naïve animals **(d)**. **(e)** *p*-values were determined by two-way ANOVA and Dunn’s multiple comparison test comparing CD45iv^−^ versus CD45iv^+^. Differences were considered significant at *p* □ 0.05 with exact *p* values displayed in the figure.

In separate experiments, serum from Wistar rats vaccinated with CV2CoV or CV2CoV.617.2 mRNA vaccines (monovalent; 8 μg), or vaccines combining CV2CoV617.2 with either CV2CoV or CV2CoV.351 (bivalent; 4 μg of each) on Day 0 and Day 21 (Fig. S5) contained high level nAb titres against Delta (Fig. 2c). Neutralizing antibody titres were notably diminished against Omicron BA.1 in all but the bivalent vaccine group containing Beta and Delta S-protein sequences. Including the Beta S-protein sequence in the vaccine resulted in nAb titres that were 2× higher against Omicron BA.1 than those induced by Delta alone, whereas the nAb titres induced by vaccines without Beta S-protein were 3–9× lower against Omicron BA.1 than those induced by Delta (Fig. 2c). The bivalent formulation contained a half dose of each mRNA compared with the monovalent vaccines with better antigen coverage; this strategy may be advantageous in case of the emergence of additional antigenic distant variants.

Cellular immunity also contributes to protection against COVID-19 and recent evidence suggests that viral-vector vaccines and mRNA vaccines elicit long-lasting S-specific CD4^+^ and CD8^+^ T cell responses with broad cross-reactivity against VOCs.^19^ While tissue resident memory T cells (T_RM_) are induced by high concentration mRNA vaccines;^20^ we investigated if lower concentrations of mono- or bivalent mRNA vaccines could induce lung parenchymal tissue resident T cells. We observed that both the monovalent and bivalent mRNA vaccines triggered potent S-specific CD4^+^ and CD8^+^ responses, with 2–3 log increases in lung CD45iv^−^ CD8^+^ T cells, compared with naïve non-vaccinated animals (Fig. 2d and Fig.S4a). Further characterization of this T cell population, using markers of lung T cell residency, confirmed significantly higher proportions of T_RM_ cells compared with the CD45iv^+^ populations (Fig. 2e and Fig.S4b-e), while mRNA vaccination also induced an enrichment of IFNγ and granzyme-B producing CD8^+^ T cells in the lung parenchyma (Fig.S4f-i).

The accumulation of memory cells at mucous membrane sites is critical for the control of viral pneumonia and their presence has been reported to be associated with less severe COVID-19 symptoms.^21^ CD8^+^ T cells may contribute to protection when antibodies titres are suboptimal in non-human primates.^22^ The role of the durability and specificity of T_RM_ CD8^+^ cells in protection from disease requires further investigation using antibody-mediated depletion. Depletion of CD8^+^ cells following immunization with mRNA vaccine, prior to SARS-CoV-2 challenge, led to different outcomes in a transgenic mouse model, possibly due to the vaccine dose used and variability in nAb concentrations.^23,24^ Interest in vaccine-elicited T cell responses has increased due to immune escape by various VOCs, including Omicron.^19^

Preservation of the epitope repertoire may be critical for the defence against current and future VOCs and may bring substantial benefits by contributing to protection against severe disease. This feature along with tissue residency makes the bivalent mRNA vaccine, and more generally, multivalent vaccines, highly appropriate candidates for further development.

In summary, SARS-CoV-2 evolution is a challenge for vaccine-based strategies for disease control. Our study demonstrates that a low-dose, bivalent, unmodified mRNA vaccine is highly efficacious in pre-clinical mouse and rat models and suggests that dose-sparing, multivalent vaccines combining mRNA encoding the S-protein from the variants with unrelated lineages may induce heterologous protection and thus increase the breadth of immune responses. Given their exceptional flexibility in antigen formulation, mRNA vaccine platforms offer advantages regarding adaptability to circulating VOCs and opportunities to design pan-sarbecovirus vaccines.

## Acknowledgements

The authors would like to thank Mareen Lange, Patrick Zitzow, and Laura Timm for their excellent technical assistance, Andrea Aebischer for protein preparations, Claudia Wylezich for sequencing, Frank Klipp, Doreen Fiedler, Christian Lipinski, Steffen Kiepert, Bärbel Berger, and Kerstin Kerstel for their invaluable support in the animal facility, Julia Führer, Julia Schröder, Nathalie Simon, Sonja Heidu, LuisaMarques Palma, Martina Schmickl for their outstanding laboratory work, Annette Moebes, Nina Schneck, Michaela Trapp, Jessica Wild, Nadine Platner and Lisa Wellhäuser for their relentless support in the lab, and Domenico Maione for critical review of the manuscript.

We would also like to thank Ronald Dijkman for providing the SARS-CoV-2 B.617.2 isolate, Adam Taylor (Alchemy Medical Writing, UK; funded by CureVac AG, Germany) and Margaret Haugh (CureVac AG) for medical writing and editorial services, and Giulia Povellato for reviewing the drafts.

## Funding

This work was funded by the German Federal Ministry of Education and Research (BMBF; grant 01KI20703), core funds of the German Federal Ministry of Food and Agriculture, and CureVac AG.

## Methods

### Ethics

The animal experiments were evaluated and approved by the ethics committee of the State Office of Agriculture, Food safety, and Fishery in Mecklenburg–Western Pomerania (LALLF M-V: LVL MV/TSD/7221.3-1-055/20) and the State Office for Occupational Safety, Consumer Protection and Health in Brandenburg (LAVG: 2347-5-2021). All procedures using SARS-CoV-2 were carried out in approved biosafety level 3 (BSL3) facilities.

### Study design

K18-hACE2 transgenic mice were vaccinated on Day 0 (prime) and Day 28 (boost) and infected (challenge) on Day 56, as detailed in Fig. S1. The animals were infected under short-term isoflurane inhalation anesthesia with 25 μl of either 10^4.4^ TCID_50_ SARS-CoV-2 lineage B.1.617.2 Delta (calculated from back-titration of the original material) or 10^4.4^ TCID_50_ SARS-CoV-2 lineage B.1.351 Beta (calculated from back-titration of the original material) per animal. An oral swab sample of each animal, under short-term isoflurane inhalation anaesthesia, was taken 4 days after infection. Animals with signs of severe clinical symptoms and/or body weight loss over 20% were euthanized immediately, all remaining animals were euthanized at Day 10 post infection.

RNA from mouse nasal swabs and organ samples was extracted using the NucleoMag^®^ VETkit (Macherey-Nagel, Düren, Germany) in combination with a Biosprint 96 platform (Qiagen, Hilden, Germany). Each extracted sample was eluted in 100 μl. Viral RNA genome was detected and quantified by real-time reverse transcription polymerase chain reaction (RT-qPCR) on a BioRad real-time CFX96 detection system (BioRad, Hercules, USA). Target sequence for amplification was the viral RNA-dependent RNA polymerase.^25^ Genome copies per μl RNA template were calculated based on a quantified standard RNA, where absolute quantification was done by the QX200 Droplet Digital PCR System in combination with the 1-Step RT-ddPCR Advanced Kit for Probes (BioRad, Hercules, USA). The limit of detection was calculated to be 1 genome copy/μl RNA. Samples (mouse swabs/organs) that tested positive for viral genomic RNA were evaluated using an assay specifically detecting sgRNA of the ORF7a as described in Hoffmann et al 2021.^26^

Wistar rats were vaccinated on Day 0 (prime) and Day 21 (booster), as detailed in Fig. S5.

### Vaccine

The mRNA vaccine is based on the RNActive^®^ platform (claimed and described in e.g. WO2002098443 and WO2012019780) and is comprised of a 5’ cap structure and untranslated region (UTR), a GC-enriched open reading frame, 3’ UTR, polyA tail and does not include chemically modified nucleosides. The mRNA was encapsulated using the lipid nanoparticle (LNP) technology of Acuitas Therapeutics (Vancouver, Canada). The LNPs used in this study are particles of ionizable amino lipid, phospholipid, cholesterol and a PEGylated lipid. The mRNA encoded protein is based on the S-protein of SARS-CoV-2 NCBI Reference Sequence NC_045512.2, GenBank accession number YP_009724390.1 and encodes for full length S featuring K986P and V987P mutations.

### SARS-CoV-2 propagation and handling

SARS-CoV-2 B.1.617.2-lineage hCoV-19/Switzerland/BE-IFIK-918-4879/2021 (GISAID accession EPI_ISL_1760647) “Delta” was kindly provided by Ronald Dijkman, Institute for Infectious Diseases, University of Bern, Switzerland. SARS-CoV-2 hCoV-19/Germany/NW-RKI-I-0029/2020 B.1.351-lineage (GISAID accession EPI_ISL_803957) “Beta” was kindly provided by Robert-Koch-Institut, Berlin, Germany. SARS-CoV-2 B.1.1.529 sublineage BA.1 “Omicron” FFM-ZAF0396/2021; GenBank accession: OM617939.1, GISAID accession EPI_ISL_6959868^27^ was used for virus neutralization assay. Virus stocks were propagated (three passages for Delta, two passages for Beta and Omicron) on Vero E6 cells (Collection of Cell Lines in Veterinary Medicine CCLV-RIE 0929) using a mixture of equal volumes of Eagle MEM (Hanks’ balanced salts solution) and Eagle MEM (Earle’s balanced salts solution) supplemented with 2 mM L-Glutamine, nonessential amino acids adjusted to 850 mg/L, NaHCO_3_, 120 mg/L sodium pyruvate, 10% fetal bovine serum (FBS), pH 7.2. The virus was harvested after 72 hours, titrated on Vero E6 cells and stored at −80 °C until further use.

### Serology/Antibody ELISA

Antibodies reactive against the receptor binding domain (RBD) of the ancestral SARS-CoV-2 were measured using the established ELISA protocol as previously described.^26,28^

To evaluate specifically the presence of virus-neutralizing antibodies in serum samples the virus neutralization test was performed with use of the viruses introduced for the challenge infection and Omicron in concentrations as described.^26^

### Flow cytometry

Vaccine-induced T cells were characterized by flow cytometry. To discriminate between parenchymal and vascular T cells, vaccinated mice received 3 μg (in PBS) of anti-mouse CD45 antibody (retro-orbital administration) for 3 minutes during lethal anesthesia. Spleen and lung tissue from vaccinated mice or naïve controls were harvested and kept at 4°C before either mechanical disruption (spleen) or mechanical disruption followed by enzymatic digestion (lung: 175 μg/ml LiberaseTM [Roche] and 0.1 mg/ml DNase I in serum-free medium) to generate single cell suspensions. Erythrocyte lysis in both suspensions was performed using 1 × red blood cell (RBC) lysis buffer (BioLegend) before determining cell counts (Biorad TC20 automated cell counter). After an additional washing step, cell surface receptor staining began with Zombie UV™ Fixable dye (1:100; BioLegend) for 20 minutes in the dark at 4°C followed by two washing steps. Unspecific antibody binding was blocked with TruStain FcX (anti-mouse CD16/32) solution (BioLegend) for 5 minutes at 4°C before adding freshly prepared antibody cocktails. Cells were incubated with surface antibodies for 20 minutes at 4°C in the dark, followed by two washing steps before fixation for 30 minutes at room temperature with 4% paraformaldehyde (PFA). SARS-CoV-2 S-peptide specific responses were investigated by culturing single cell suspensions in the presence of 0.5 μg/ ml PepMix™ SARS-CoV-2 (JPT) for 15 hours and brefeldin A (BioLegend) for an additional 4 hours. For intracellular antibody staining cells were washed twice before fixated with intracellular fixation buffer (eBioscience, Foxp3/Transcription Factor Staining Buffer Set) and washed with 1 × permeabilization buffer followed by incubation of antibody cocktails diluted in 1 × permeabilization buffer. Details of the antibody clones, fluorochromes, dilutions, and fluorescent minus one (FMO) controls are provided in Table S4. Cells were stored in PBS at 4°C in the dark for no longer than 24 hours before acquisition on a BD Fortessa™ instrument. Details of the T cell gating strategy are presented in Fig. S3.

## Notes

### Competing Interest Statement

B Corleis, A Dorhoi, B Petsch and M Beer declare institutional funding for the work. SO Mueller and B Petsch declare holding company shares or stock options. The remaining authors declare no competing interests.

