## Supplemental Fig. S1-S5 and Table S1-S4 for "Low-dose bivalent mRNA vaccine is highly effective against different SARS-CoV-2 variants in a transgenic mouse model"

### Supplemental Tables and Figures

#### Fig. S1. Experimental design (mouse challenge experiments)

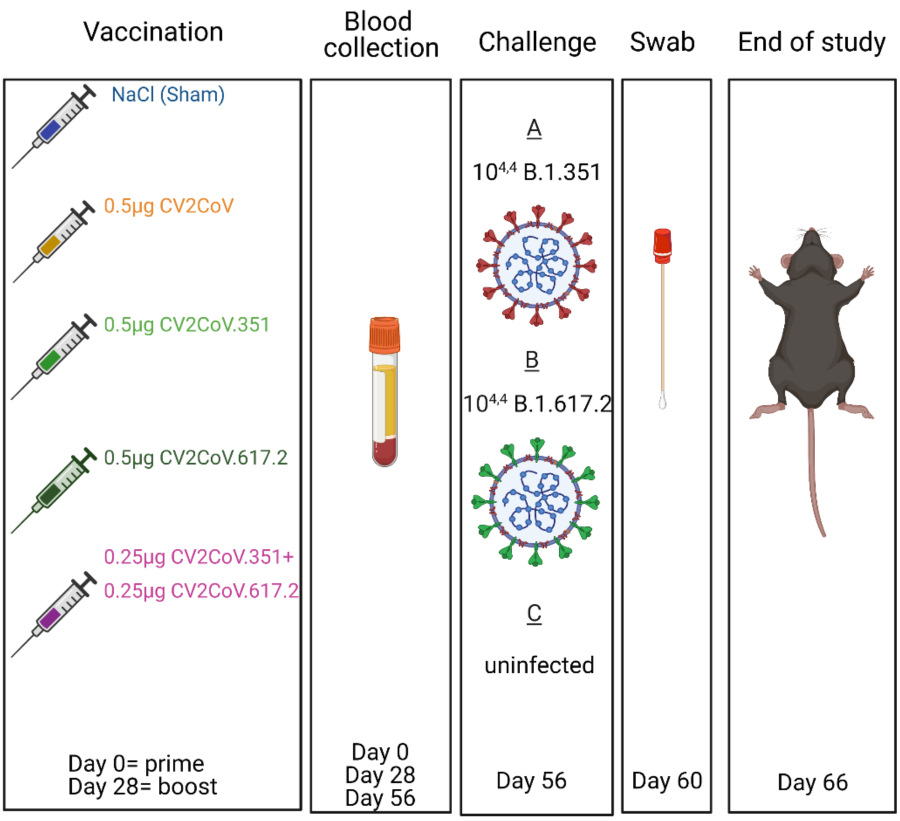


K18-hACE2 mice were vaccinated on Day 0 (prime) and Day 28 (boost) intramuscularly with 20 µl of either 0.5 µg CV2CoV (ancestral), 0.5 µg CV2CoV.351 (Beta), 0.5 µg CV2CoV.617.2 (Delta), 0.25 µg CV2CoV.351 + 0.25µg CV2CoV.617.2 (Beta and Delta) or received 20 µl NaCl (sham) as a control. Blood samples for analysis of humoral responses were collected at Day 0, 28 and 56. For the mRNA vaccinated groups, on Day 56, 4 mice per group were euthanized before viral challenge to analyse the vaccine induced cellular immune response in lung and spleen. The remaining mice were challenged with either $10^{4.4}$ B.1.351 or $10^{4.4}$ SARS-CoV-2 B.1.617.2. At 4 days post-challenge an oral swab was taken from all mice. At 10 days post-challenge, or when animals became too sick (humane endpoint), the animals were euthanized, and organs (lung, spleen, conchae, cerebrum, and cerebellum) were taken for analysis of viral load and cellular immune responses. The numbers of mice per assay are summarized in Table S1. Image was generated using the illustration software Biorender (Biorender.com).

**Fig. S2.** **Protection against challenge with SARS-CoV-2 B.1.351 (Beta) or B.1.617.2 (Delta) with monovalent and bivalent mRNA vaccines.**

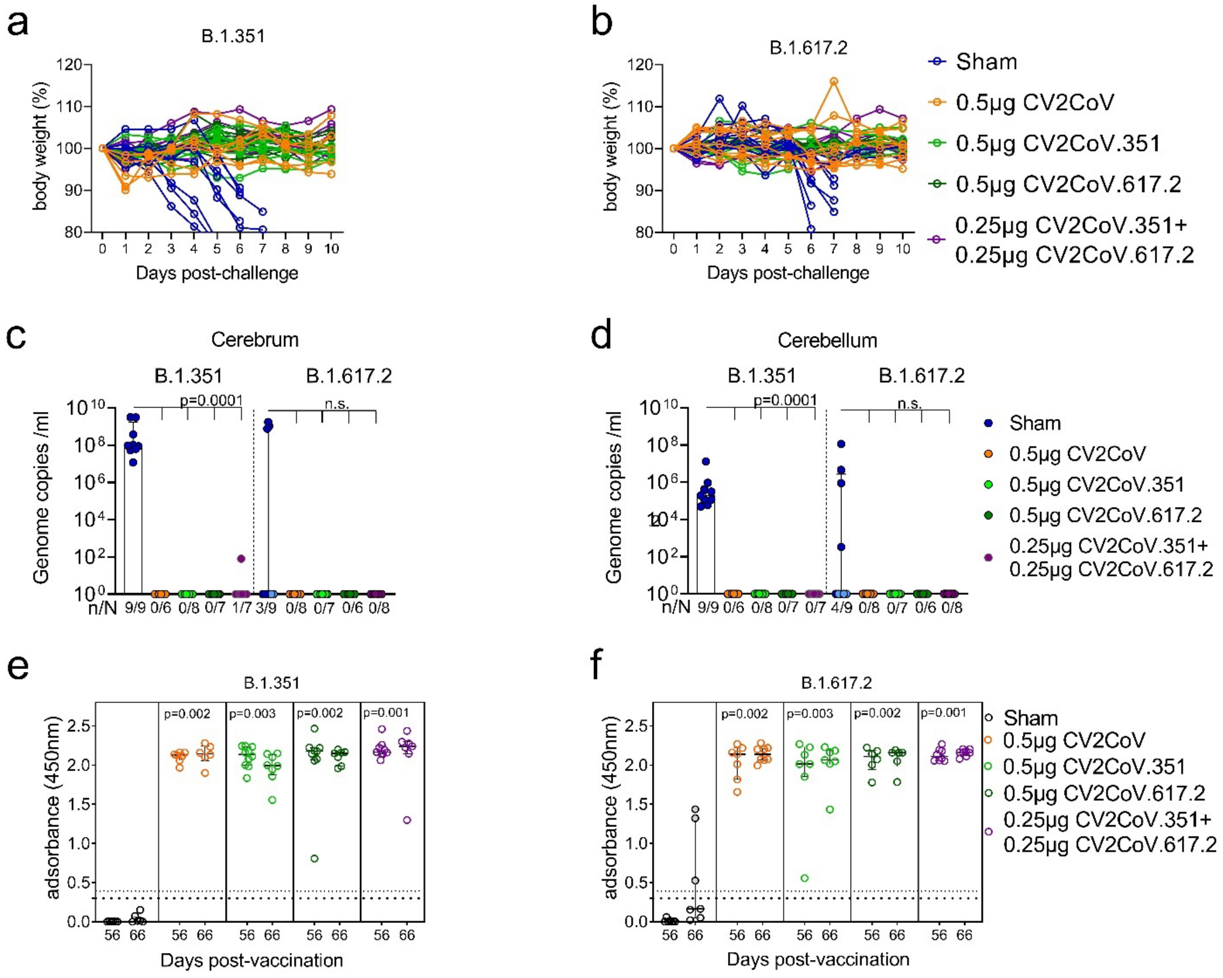


Vaccinated K18-hACE2 mice were challenged with SARS-CoV-2 variants as described in Fig. S1. The percentage change in body weight of mice infected with SARS-CoV-2 B.1.351 **(a)** or B.1.617.2 **(b)** was monitored daily. Lines represent individual animals over the course of the experiment. RT-qPCR results from Day 10 cerebrum **(c)** and cerebellum **(d)**. Sera from all vaccinated and sham animals either before challenge (Day 56) or after challenge with B.1.351 **(e)** or B.1.617.2 **(f)** (Day 66) were analysed in an RBD ELISA (ancestral RBD) for total Ig anti-RBD antibodies. Sham group samples were obtained at Day 10 (filled circle) or at the humane endpoint (transparent circle). *p* Values were determined by one-way ANOVA and Dunn's multiple comparison test against the sham group (**c-f**).

**Fig. S3. T-cell gating strategy**

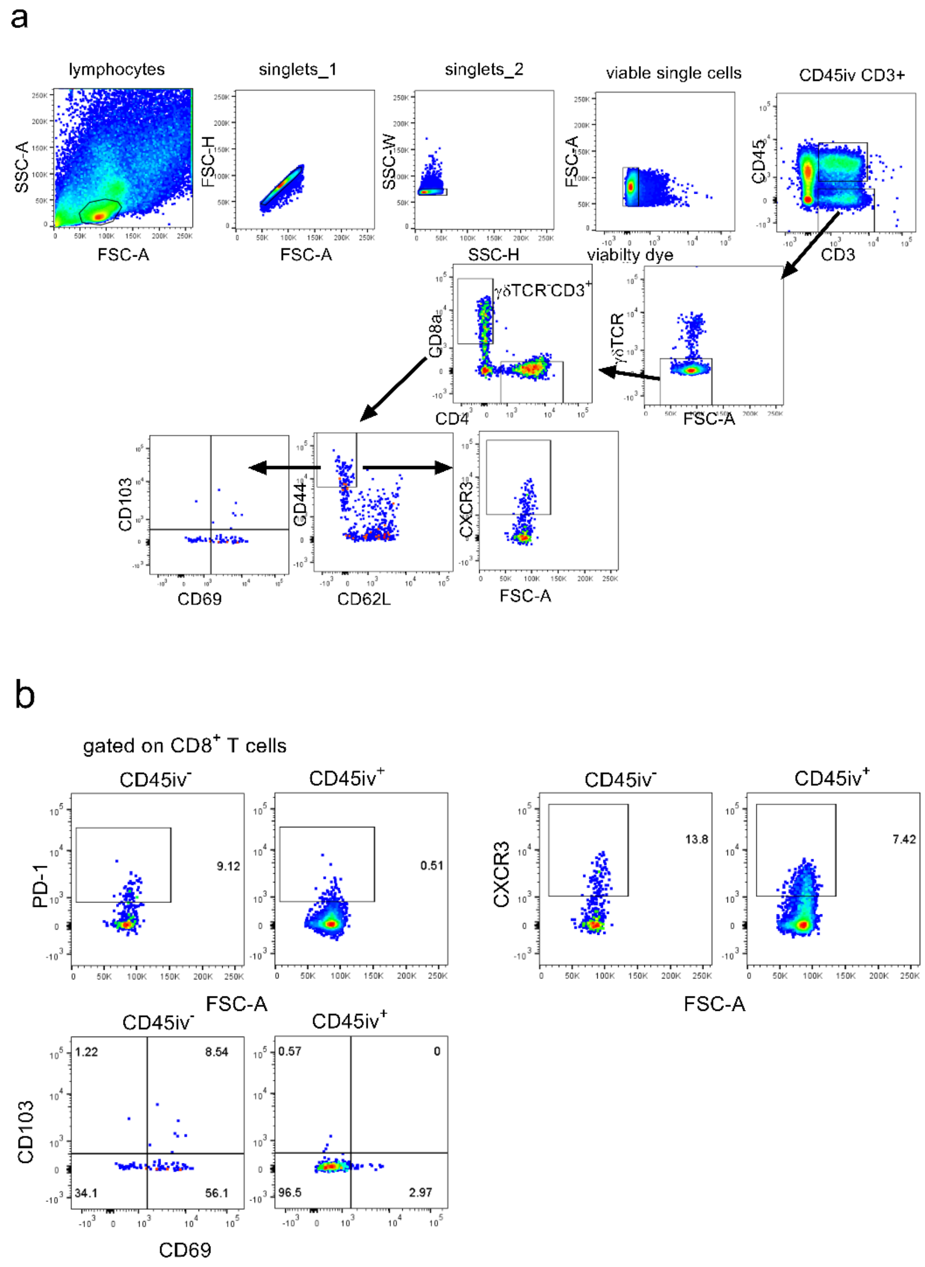


Gating strategy for T cell analysis **(a)**. Lymphocytes were identified by SSC-A vs FSC-A and lymphocyte doublets were excluded by FSC-H/FSC-A and SSC-W/SSC-H. Dead cells were eliminated using a fixable live/dead viability dye (UV zombie). To distinguish lung parenchymal (CD45iv$^-$) from vascular (CD45iv$^+$) CD3$^+$ T cells, 3µg anti-mouse CD45 antibody was injected (i.v.) for 3 minutes during lethal anesthesia. CD3$^+$ T cells were further analysed by exclusion of γδTCR$^+$ cells before gating on CD8$^+$ cytotoxic T cells and CD4$^+$ helper T cells. CD4$^+$ and CD8$^+$ T cell subsets were further analysed for the frequency of CXCR3$^+$ T cells or T$_{RM}$ cells defined as CD45iv$^-$CD3$^+$γδTCR$^-$ CD8$^+$CD44highCD62L$^-$CD103$^+$CD69$^+$. Flow plots are generated from one representative mouse sample.

Gating strategy CD45iv$^-$ versus CD45iv$^+$ T cell analysis **(b)**. T cells were analysed as described above **(a)**. CD45iv$^-$ and CD45iv$^+$ T cells were compared for their expression of markers associated with tissue residency and lung migration such as PD-1, CXCR3, CD103 and CD69. Flow plots are generated from one representative mouse sample.

**Fig. S4.** **Induction of tissue resident T cells and S-peptide specific responses by monovalent and bivalent mRNA vaccines.**

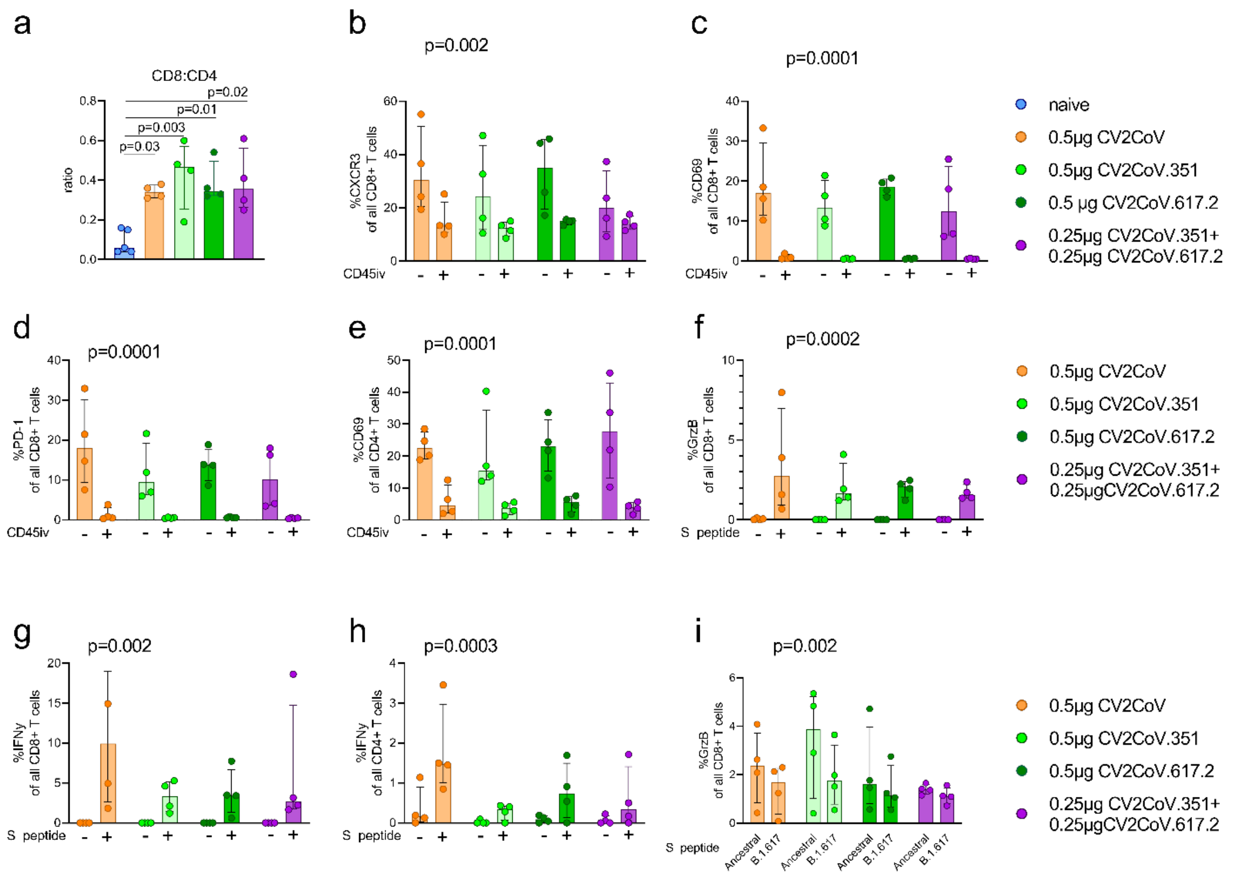


Naïve mice served as controls for analysis of T cell responses induced by monovalent and bivalent mRNA vaccines. At Day 56 post-first-dose, lung parenchyma T cells were analysed by *in vivo* injection of 3 µg anti-mouse CD45 antibodies (CD45iv) for 3 minutes before harvesting of lung tissue. **(a)** CD8:CD4 ratio of CD45iv$^-$CD3$^+$ T cells in lung tissue. **(b-e)** Frequency of markers associated with tissue resident ($T_{RM}$) cells on CD45iv$^-$ versus CD45iv$^+$CD8$^+$ **(c-d)** or CD4$^+$ **(e)** T cells. Granzyme B production by lung CD8$^+$ T cells **(f)** and IFNy production by CD8$^+$ T cells **(g)** or CD4$^+$ T cells **(h)** was investigated by *in vitro* re-stimulation of lung cells with S-peptide pools derived from ancestral SARS-CoV-2.

**(i)** Comparison of Granzyme B production by lung CD8$^+$ T cells after stimulation with S-peptide pools derived from ancestral SARS-CoV-2 (ancestral) or B.1.617.2 (Delta). Each dot represents one individual mouse.

Scatter plots are labelled with median and interquartile range. *p v*alues were determined by one-way ANOVA and Dunn's multiple comparison test against the naïve group **(a)** or two-way ANOVA and Dunn's multiple comparison test comparing CD45iv$^-$ versus CD45iv$^+$ **(b-e)** or unstimulated (-) versus S-peptide stimulated (+) conditions **(f and i)**. Differences were considered significant at $p < 0.05$ with exact p values displayed in the figure.

**Fig. S5.** **Experimental Design (rat experiment)**

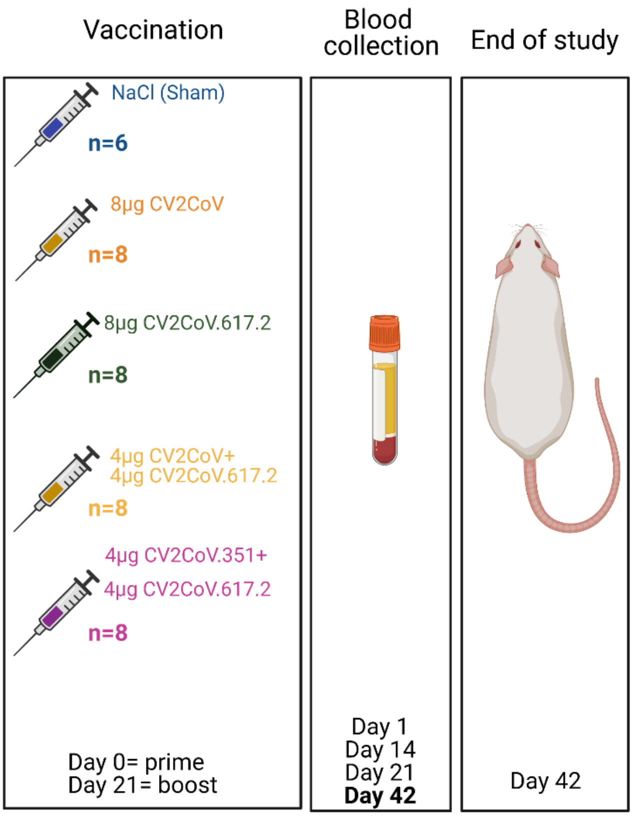


Wister rats were vaccinated on Day 0 (prime) and Day 21 (boost) intramuscularly with either 100 µl of 8 µg CV2CoV (ancestral), 8 µg CV2CoV.617 (Delta), 4 µg CV2CoV + 4 µg CV2CoV.617 (ancestral and Delta), 4 µg CV2CoV.351 + 4 µg CV2CoV.617.2 (Beta and Delta) mRNA vaccine or received 100 µl NaCl (sham) as a control group. Blood samples were collected at Day 1, 14, 21 and Day 42. Cross-response to Omicron and Delta were only tested using Day 42 samples. Image was generated using the illustration software Biorender (Biorender.com).

**Table S1. Numbers of mice for per assay**

| Group | Total (n) | Survival data (n) | Viral loads (n) | nAb titres pre-challenge (n) | nAb titres post-challenge (n) | T cells pre-challenge (n) | RBD Elisa pre-challenge (n) | RBD Elisa post-challenge (n) |
|---|---|---|---|---|---|---|---|---|
| **Experiment 1 (Challenge virus: SARS-CoV-2 Beta)** | | | | | | | | |
| Sham | 9 | 9 | 9 | N/A | N/A | N/A | 9 | 8 |
| CV2CoV | 8 | 6 | 6 | 8 | 5 | 2 | 7 | 6 |
| CV2CoV.351 (Beta) | 10 | 8 | 8 | 10 | 8 | 2 | 10 | 8 |
| CV2CoV.617.2 (Delta) | 9 | 7 | 7 | 9 | 7 | 2 | 9 | 7 |
| CV2CoV.351+CV2CoV.617.2 | 9 | 7 | 7 | 9 | 7 | 2 | 9 | 7 |
| **Experiment 2 (Challenge virus: SARS-CoV-2 Delta)** | | | | | | | | |
| Sham | 9 | 9 | 9 | N/A | N/A | N/A | 9 | 7 |
| CV2CoV | 10 | 8 | 8 | 10 | 8 | 2 | 8 | 8 |
| CV2CoV.351 (Beta) | 9 | 7 | 7 | 9 | 7 | 2 | 7 | 7 |
| CV2CoV.617.2 (Delta) | 8 | 6 | 6 | 8 | 6 | 2 | 6 | 6 |
| CV2CoV.351+CV2CoV.617.2 | 10 | 8 | 8 | 10 | 8 | 2 | 8 | 8 |
| **Naive animals** | | | | | | | | |
| Non-vaccinated or challenged | N/A | N/A | N/A | N/A | N/A | 5 | N/A | N/A |

Abbreviations: N/A, not applicable; RBD, receptor-binding domain; nAB, neutralizing antibody.

**Table S2. Viral RNA genome load detected in oral swab samples Day 4 post-challenge) expressed as Cq value for total RNA and subgenomic RNA (ORF7a)**

| Vaccine | Challenge | Mouse ID | Genomic RNA | Subgenomic RNA |
|---|---|---|---|---|
| CV2CoV | Delta B.1.617.2 | 25.2.3 | 37.6 | nd |
| none | Beta B.1.351 | 24.3.3 | 35.0 | nd |
| | | 24.3.4 | 29.8 | 36.1 |
| | | 24.3.5 | 33.0 | nd |
| | | 24.4.1 | 36.0 | nd |
| | | 24.4.2 | 34.3 | nd |
| | | 24.4.3 | 34.5 | nd |
| | | 24.4.4 | 31.7 | nd |
| | | 24.4.5 | 36.5 | nd |
| | Delta B.1.617.2 | 25.3.1 | 38.0 | nd |
| | | 25.3.2 | 38.1 | nd |
| | | 25.3.3 | 35.4 | nd |
| | | 25.4.2 | 36.3 | nd |
| | | 25.4.4 | 36.8 | nd |
| | | 25.4.5 | 37.8 | nd |

Abbreviation: Cq, RT-PCR cycle number at which reaction curves intersected the threshold line (also referred to as the crossing point); CV2CoV, ancestral mRNA vaccine; nd, not detected.

This table includes data for Day 4 swab samples that were positive for SARS-CoV-2 genomic RNA (Cq <45); all other Day 4 swab samples were negative (Cq ≥45). Subgenomic RNA analysis was only conducted for Day 4 swab samples that tested positive for genomic RNA.

**Table S3 Viral RNA genome load detected in organ samples (10 dpc or indicated), expressed as Cq value for total RNA and subgenomic RNA (ORF7a)**

| Vaccine | Challenge | Mouse ID (days post-challenge) | Organ | Genomic RNA | Subgenomic RNA |
|---|---|---|---|---|---|
| none | Beta B.1.351 | 24.3.2 (7) | Cerebellum | 21.6 | 23.1 |
| | | | Cerebrum | 14.1 | 15.4 |
| | | | Conchae | 35.5 | nd |
| | | | Lung | 30.0 | 33.2 |
| | | 24.3.3 (6) | Cerebellum | 26.5 | 28.6 |
| | | | Cerebrum | 19.0 | 20.6 |
| | | | Conchae | 30.4 | 35.4 |
| | | 24.3.4 (5) | Cerebellum | 25.3 | 25.7 |
| | | | Cerebrum | 17.0 | 18.2 |
| | | | Conchae | 26.3 | 29.7 |
| | | | Lung | 19.4 | 22.2 |
| | | 24.3.5 (5) | Cerebellum | 29.1 | 30.7 |
| | | | Cerebrum | 19.7 | 21.4 |
| | | | Conchae | 28.2 | 33.6 |
| | | | Lung | 17.4 | 20.4 |
| | | 24.4.1 (7) | Cerebellum | 28.2 | 31.5 |
| | | | Cerebrum | 14.1 | 17.1 |
| | | | Conchae | 30.9 | 35.9 |
| | | | Lung | 38.4 | 28.4 |
| | | 24.4.2 (6) | Cerebellum | 28.1 | 30.0 |
| | | | Cerebrum | 18.8 | 20.3 |
| | | | Conchae | 27.5 | 31.8 |
| | | | Lung | 28.2 | 31.2 |
| | | 24.4.3 (5) | Cerebellum | 26.8 | 28.4 |
| | | | Cerebrum | 19.5 | 21.1 |
| | | | Conchae | 25.2 | 29.0 |
| | | | Lung | 19.6 | 21.8 |
| | | 24.4.4 (5) | Cerebellum | 29.3 | 30.9 |
| | | | Cerebrum | 21.8 | 22.5 |
| | | | Conchae | 29.1 | 34.8 |
| | | | Lung | 26.3 | 29.2 |
| | | 24.4.5 (6) | Cerebellum | 27.5 | 28.6 |
| | | | Cerebrum | 18.9 | 20.5 |
| | | | Conchae | 31.1 | 35.5 |
| CV2CoV | Beta B.1.351 | 24.1.3 (10) | Conchae | 36.5 | nd |
| | | 24.1.4 (10) | Conchae | 33.0 | nd |
| | | 24.1.5 (10) | Conchae | 32.0 | nd |
| | | 24.2.3 (10) | Conchae | 32.3 | nd |
| CV2CoV.351 | Beta B.1.351 | 24.5.1 (10) | Conchae | 34.1 | nd |
| | | 24.5.2 (10) | Conchae | 41.0 | nd |
| | | 24.5.4 (10) | Conchae | 35.0 | nd |
| | | 24.6.1 (10) | Conchae | 37.3 | nd |
| | | 24.6.3 (10) | Conchae | 34.7 | nd |
| | | 24.6.5 (10) | Conchae | 36.8 | nd |
| CV2CoV.617.2 | Beta B.1.351 | 24.7.2 (10) | Conchae | 30.8 | 40.9 |
| | | 24.7.3 (10) | Conchae | 36.7 | nd |
| | | 24.7.4 (10) | Conchae | 34.3 | nd |
| | | 24.7.5 (10) | Conchae | 38.4 | nd |
| | | 24.8.1 (10) | Conchae | 35.6 | nd |
| | | 24.8.3 (10) | Conchae | 35.5 | 40.8 |
| | | 24.8.5 (10) | Conchae | 38.1 | nd |
| CV2CoV.351/ CV2CoV.617.2 | Beta B.1.351 | 24.9.4 (10) | Conchae | 38.1 | nd |
| | | 24.9.5 (10) | Conchae | 38.4 | nd |
| | | 24.10.1 (10) | Conchae | 33.0 | nd |
| | | 24.10.2 (10) | Cerebrum | 37.7 | nd |
| | | | Conchae | 36.4 | nd |
| | | 24.10.3 (10) | Conchae | 36.9 | nd |

| Vaccine | Challenge | Mouse ID (days post-challenge) | Organ | Genomic RNA | Subgenomic RNA |
|---|---|---|---|---|---|
| | | 24.10.5 (10) | Conchae | 35.2 | nd |
| none | Delta B.1.617.2 | 25.3.1 (7) | Cerebellum | 22.8 | 25.2 |
| | | | Cerebrum | 14.9 | 18.9 |
| | | | Conchae | 30.4 | 34.5 |
| | | | Lung | 36.3 | nd |
| | | 25.3.2 (7) | Cerebellum | 25.2 | 28.0 |
| | | | Cerebrum | 14.2 | 16.8 |
| | | | Conchae | 27.6 | 32.1 |
| | | 25.3.3 (7) | Conchae | 29.2 | 37.2 |
| | | | Lung | 41.6 | nd |
| | | 25.3.4 (7) | Conchae | 31.3 | nd |
| | | 25.4.1 (7) | Cerebellum | 18.1 | 21.0 |
| | | | Cerebrum | 15.4 | 18.3 |
| | | | Conchae | 28.6 | 33.2 |
| | | 25.4.2 (10) | Conchae | 31.6 | 41.2 |
| | | | Lung | 33.7 | nd |
| | | 25.4.3 (10) | Conchae | 30.2 | 35.6 |
| | | 25.4.4 (10) | Conchae | 31.3 | 37.4 |
| | | | Lung | 29.4 | 31.9 |
| | | 25.4.5 (7) | Cerebellum | 36.6 | nd |
| | | | Conchae | 28.5 | 32.4 |
| | | | Lung | 26.2 | 28.9 |
| CV2CoV | Delta B.1.617.2 | 25.1.4 (10) | Conchae | 36.6 | nd |
| | | 25.2.3 (10) | Conchae | 37.5 | nd |
| | | 25.2.5 (10) | Conchae | 36.6 | nd |
| CV2CoV.351 | Delta B.1.617.2 | 25.5.1 (10) | Conchae | 36.0 | nd |
| | | 25.5.4 (10) | Conchae | 38.1 | nd |
| | | 25.6.4 (10) | Conchae | 37.9 | nd |
| CV2CoV.351/ CV2CoV.617.2 | Delta B.1.617.2 | 25.10.3 (10) | Conchae | 38.9 | nd |

Abbreviations: Cq, RT-PCR cycle number at which reaction curves intersected the threshold line (also referred to as the crossing point); nd, not detected.

This table includes data for organ samples that were positive for SARS-CoV-2 genomic RNA (Cq <45); all other organ samples were negative (Cq ≥45). Subgenomic RNA analysis was only conducted for organ samples that tested positive for genomic RNA.

**Table S4** **Antibody panels for T cell surface receptor and intracellular staining analyses in Fig. 2 and Fig. S3-4**

| Molecule | Fluorochrome | Isotype | Clone | Company | Cat # | Dilution factor |
|---|---|---|---|---|---|---|
| **Panel for T cell surface receptor analysis** | | | | | | |
| CD45(iv) | Alexa Fluor® 700 | Rat IgG2b, κ | 30-F11 | BioLegend GmbH | 103127 | 3 µg |
| CD3 | APC/Cyanine7 | Rat IgG2b, κ | 17A2 | BioLegend GmbH | 100221 | 100 |
| CD4 | Brilliant Violet 650™ | Rat IgG2a, κ | RM4-5 | BioLegend GmbH | 100545 | 150 |
| CD8a | Brilliant Violet 785™ | Rat IgG2a, κ | 53-6.7 | BioLegend GmbH | 100749 | 100 |
| TCR γ/δ | Brilliant Violet 510™ | Armenian Hamster IgG | GL3 | BioLegend GmbH | 118131 | 50 |
| CD62L | Brilliant Violet 605™ | Rat IgG2a, κ | MEL-14 | BioLegend GmbH | 104437 | 100 |
| CD103 | Brilliant Violet 711™ | Armenian Hamster IgG | 2E7 | BioLegend GmbH | 121435 | 100 |
| CD95 | APC | Rat IgG1, κ | SA367H8 | BioLegend GmbH | 152603 | 150 |
| CD44 | PE | Rat IgG2b, κ | IM7 | BioLegend GmbH | 103023 | 150 |
| KLRG1 | Brilliant Violet 421™ | Syrian Hamster IgG | 2F1/KLRG1 | BioLegend GmbH | 138413 | 100 |
| CD183/ CXCR3 | PE/ Cy7 | Armenian Hamster IgG | CXCR3-173 | BioLegend GmbH | 126515 | 100 |
| CD69 | FITC | Armenian Hamster IgG | H1.2F3 | BioLegend GmbH | 104505 | 100 |
| PD-1 | PE-Dazzle 594 | Rat IgG2b, κ | RMP1-30 | BioLegend GmbH | 109115 | 100 |
| **Panel for intracellular staining of T cells after stimulation** | | | | | | |
| CD3 | APC/Cyanine7 | Rat IgG2b, κ | 17A2 | BioLegend GmbH | 100221 | 100 |
| CD4 | FITC | Rat IgG2a, κ | RM4-5 | BioLegend GmbH | 100509 | 100 |
| CD8a | Brilliant Violet 785™ | Rat IgG2a, κ | 53-6.7 | BioLegend GmbH | 100749 | 100 |
| TCR γ/δ | Brilliant Violet 510™ | Armenian Hamster IgG | GL3 | BioLegend GmbH | 118131 | 50 |
| T-bet | Brilliant Violet 711™ | Mouse IgG1, κ | 4B10 | BioLegend GmbH | 644819 | 100 |
| RORγT | PE-CF594 | Mouse IgG2a | Q31-378 | BD | 562684 | 100 |
| FoxP3 | PE-Cy5.5 | Rat / IgG2a, κ | FJK-16s | ThermoFisher Scientific | 35-5773-80 | 100 |
| IFN-γ | Brilliant Violet 605™ | Rat IgG1, κ | XMG1.2 | BioLegend GmbH | 505839 | 150 |
| IL-17A | Brilliant Violet 421™ | Rat IgG1, κ | TC11-18H10.1 | BioLegend GmbH | 506925 | 100 |
| IL-10 | PE-Cy7 | Rat IgG2b, κ | JES516E3 | BioLegend GmbH | 505025 | 50 |
| Granzyme B | Alexa Fluor® 647 | Mouse IgG1, κ | GB11 | BioLegend GmbH | 515405 | 100 |
